# Effects of trypsin on aggregation, disaggregation and aggregate morphology of red blood cells in autologous plasma and serum

**DOI:** 10.1101/2021.06.24.449744

**Authors:** Yury A. Sheremet’ev

**Affiliations:** FSBEI HE «Privolzhsky Research Medical University» of the Ministry of Health of the Russian Federation, 10,b.1, Minin and Pozharsky Sq., 603005, Nizhny Novgorod, Russian Federation

**Keywords:** Erythrocyte aggregation, Disaggregation, Aggregation morphology, Autologous plasma and serum, Trypsin

## Abstract

We study the influence of trypsin on aggregation, disaggregation, and aggregate morphology of RBCs in autologous plasma and serum. The effect of trypsin on the surface charge of red blood cells and the aggregation of glutaraldehyde fixed cells after treatment with the enzyme was also studied. RBC aggregation was studied by means of an aggregometer and microscopic observations. The results obtained in this study indicate that trypsin treatment increases RBCs aggregation in autologous plasma and serum. The disaggregation of erythrocytes after trypsin treatment considerably decreased in autologous plasma and serum. Increase in the strength of red blood cell aggregates was observed in autologous plasma and serum. The microscopic images of RBCs aggregates indicate the formation of globular (pathologic) structures of aggregates in autologous plasma and serum. Trypsin decrease the surface charge of RBCs. In autologous plasma and serum, the cup shapes of RBCs appear. The control RBCs fixed with glutaraldehyde were not aggregated after their placement in autologous plasma. At the same time, red blood cells pretreated with trypsin and fixed with glutaraldehyde interact with each other in autologous plasma. The physiological significance of glycoproteins of erythrocyte surface for RBCs aggregation was discussed.

## 1. Introduction

The reversible aggregation of red blood cells (RBCs) is crucial for regulating blood viscosity changes at low shear stress [1].

Human RBCs aggregation is determined by two types of biophysical and physicochemical factors: suspending phase properties and RBC properties [2,3]. Plasma proteins, such as fibrinogen, α_2_-macroglobulin, and immunoglobulins play a crucial role in RBC aggregation [4–8]

Among the cellular factors of RBC aggregation, an important role belongs to the change in surface charge, deformability and shape [3, 9–11].

The treatment of human red blood cells with the enzyme trypsin could lead to increase their aggregation in autologous plasma, solutions of fibrinogen and high-molecular dextran [3, 12–16]. The microscopic images of RBC aggregates indicate globular (pathologic) structure aggregates [16].

Bovine RBCs do not form any aggregate under physiologic conditions [17]. The modification of the cell surface by trypsin induced rouleaux formation, whereas the modification of the cell surface by neuraminidase did not cause any aggregate formation [18].

The mechanisms of action of trypsin on the aggregation of RBCs remains unclear.

The purpose of the present investigation was to study the effects of trypsin on aggregation, disaggregation (aggregate strength), and aggregate morphology of RBCs in autologous plasma and serum. To study also the effect of trypsin on the surface charge of RBCs and the aggregation of the cells fixed with glutaraldehyde after treatment with their enzymes. RBCs aggregation was studied by means of an aggregometer and microscopic observations.

## 2. Materials and methods

### 2.1. Materials

Trypsin, NaCI, tris buffer, phosphate buffer were purchased from Sigma Chemical Co, St. Louis, MO, USA.

### 2.2. Preparation of RBCs

Venous blood was collected into vacuum tubes containing 3.8% sodium citrate (9:1). Immediately after collection, citrated blood was centrifuged at 1400 x g for 20 min, the plasma was aspirated and saved, and the buffy coat was discarded. RBCs were then washed three times with an excess volume of 150 mM NaCI saline. To obtain serum, blood was collected into vacuum tubes which did not contain anticoagulants.

### 2.3. Treatment with trypsin

The washed RBCs were preincubated in trypsin solution (10 mM Tris-HCI, 150 mM NaCI, pH 7.4). Trypsin was used at 5 mg enzyme per ml of cells [3]. Treatment with trypsin involved a 30 min incubation of RBCs in the trypsin solution (at hematocrit 10% and at 370C), and then three washes in 10 volumes of buffer. In some experiments, the intact and trypsin treatment RBCs were fixed in a 0.25% solution of glutaraldehyde prepared in phosphate buffer (10 mM Na_2_HPO_4_, 150 mM NaCI, pH 7.4) and then three washed in 10 volumes of tris buffer. After that, the RBCs were placed in the autologous plasma.

### 2.4. Assessment of RBCs aggregation

RBCs aggregation was investigated using a rheoscope designed by Schmid-Schönbein et al. [19] in our modification [11,20]. For investigation of aggregation, RBCs were resuspended in plasma or serum, and the hematocrit was adjusted to 35±0.5%. To evaluate the process of aggregation, we used the following parameters:

1. Extent of RBCs aggregation (Ma) – maximal amplitude of the aggregogram (mm).
2. Half time (t _1/2_) – half time of kinetics of aggregation (sec).
3. Extent of disaggregation – ratio (%) between disaggregation amplitude to Ma at the shear rates 10 s^−1^, 15s^−1^, 20 s^−1^ – D _10_, D _15_, D _20_.

### 2.5. Aggregate morphology

The morphology of the RBCs aggregates was studied by an optical microscope (Primo Star, Carl Zeiss, Germany). For this purpose, plasma or serum was mixed with RBCs in a ratio of 2:1. A drop of the obtained mixture of plasma or serum with RBCs was placed on a glass slide and stirred. An objective with a magnification of 100^x^ was immersed into the mixture. The total magnification of the microscope was 1000x [21].

### 2.6. Electrophoretic measurement

RBC electrophoretic mobility was determined as described by Seaman et al. [22], using a system consisting of a DC power supply, a precision glass capillary sample tube fitted with Ag/AgCI electrodes, and a microscope with a 40x water immersion objective.

### 2.7. Statistical analysis

Data are presented as the mean ± standard error mean (SEM). The results of this study were tested by using nonparametric statistics methods using the Mann-Whitney test. The level of significance was considered as ρ value <0.05.

## 3. Results

It was shown that trypsin increases RBCs of aggregation in autologous plasma (Table 1). Thus, the degree of aggregation increased by 16% and t_1/2_ reduced by more than 50%. Characteristic morphological images corresponding to RBC samples in autologous plasma from control (1A) and after trypsin treatment (1B). In contrast to the roleaux formed in the control, globular RBC aggregates were observed after trypsin treatment. Our data are consistent with previous reports [16].

**Table 1.** Effect of trypsin on RBCs aggregation

| Aggregation<br>indexes | RBCs + plasma |  | RBCs + serum |  |
| --- | --- | --- | --- | --- |
|  | Control | Trypsin | Control | Trypsin |
| Ma (mm) | 85.42±1.44 | 99.56±3.05* | 63.21±1.93 | 93.82±3.1* |
| t<sub>1/2</sub> (sec) | 31.14±2.30 | 17.33±1.11* | 74.29±6.41 | 17.64±2.1* |

\*Significant difference ( $p < 0.05$ ) as compared to control

The results of the study of RBC aggregation in autologous serum are shown in Table 1. It was found out that trypsin increases RBC aggregation in autologous serum. The degree of aggregation increased, while t_1/2_ decreased. Characteristic morphological images corresponding to RBC samples in autologous serum from control (2A) and treatment with trypsin (2B) are shown in Fig.2. Fig. 2A shows the formation of rouleaux in autologous serum in control. The formation of pathologic globular structures of RBC aggregates was observed in autologous serum after treatment with trypsin (Fig. 2B). As shown in Figures 1B and 2B, after treatment of RBC with trypsin, the cup shape of erythrocytes appeared in plasma and serum.

**Fig.1.**
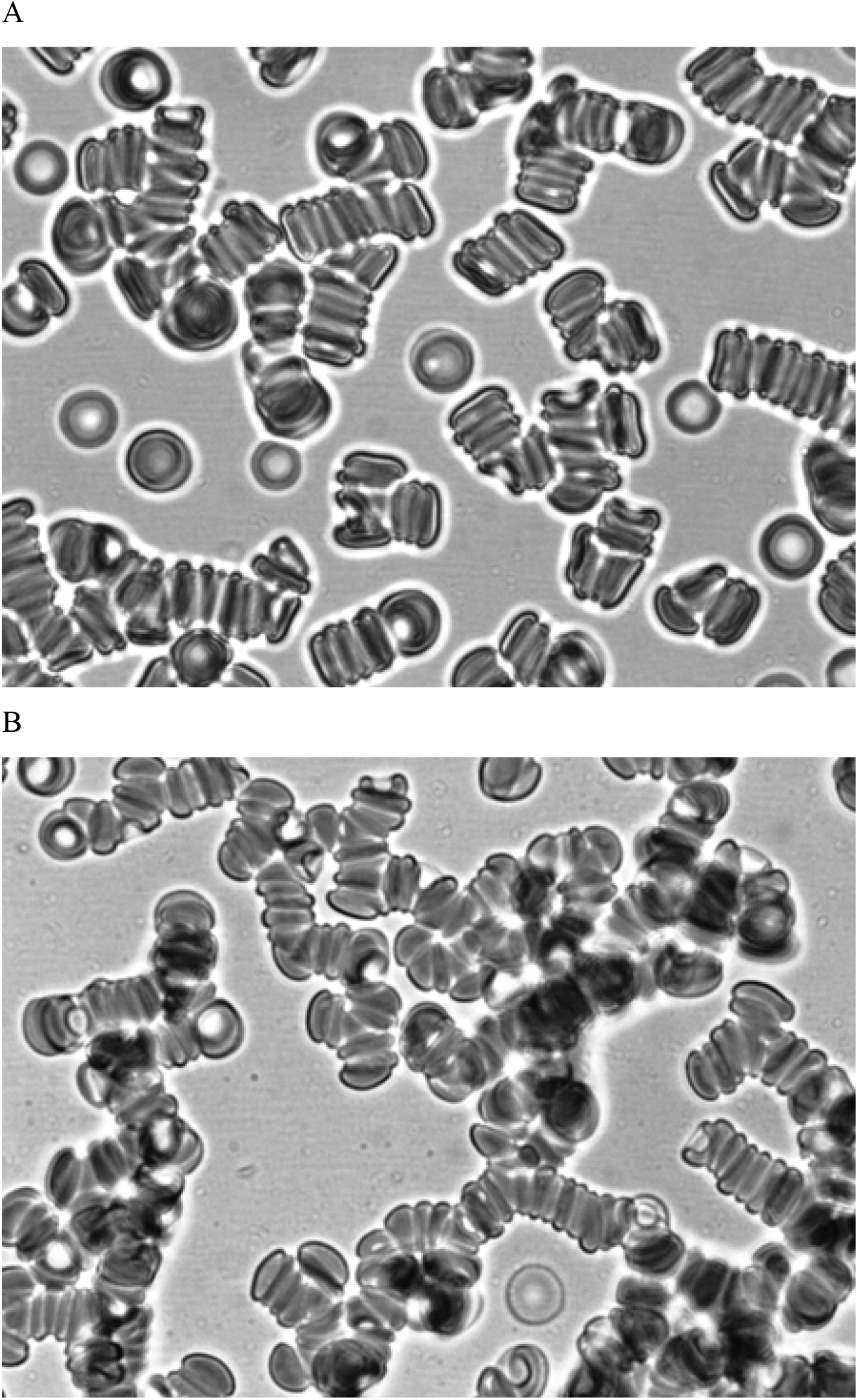
Aggregation of RBCs: control (A) and treated with trypsin (B) in autologous plasma. Magnification 1000x.

**Fig.2.**
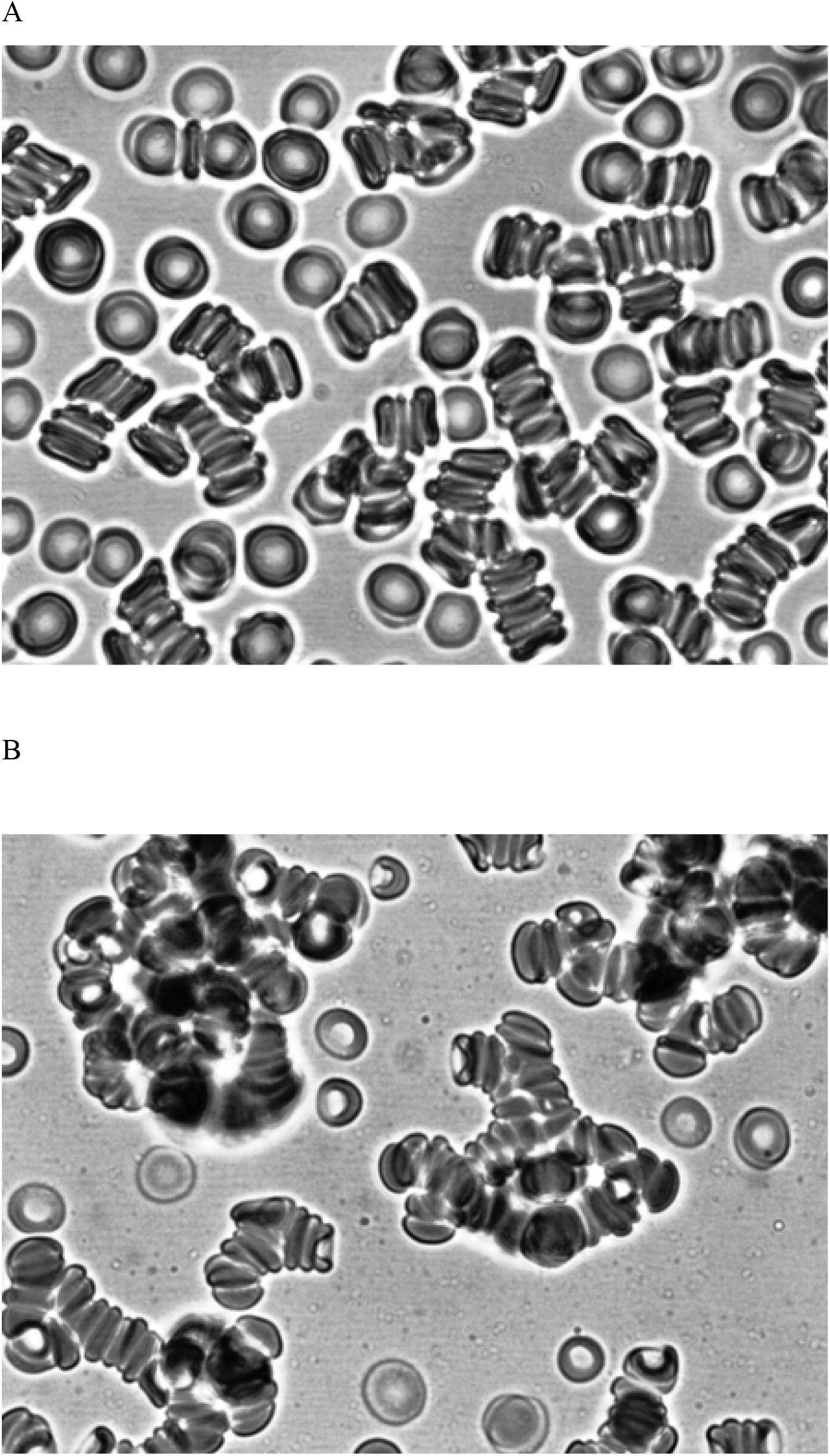
Aggregation of RBCs: control (A) and treated with trypsin (B) in autologous serum. Magnification 1000x.

Results of the study of disaggregation of erythrocytes in autologous serum are shown in Table 2. Disaggregation of erythrocytes after treatment with trypsin in autologous plasma compared to control decreased. Disaggregation of RBCs in autologous serum after treatment with trypsin also decreased at all shear rates (Table 2).

**Table 2.** Effect of trypsin on RBCs disaggregation

| Disaggregation<br>indexes | RBCs + plasma |  | RBCs + serum |  |
| --- | --- | --- | --- | --- |
|  | Control | Trypsin | Control | Trypsin |
| D<sub>10</sub> | 51.09±2.78 | 22.34±4.94 * | 60.43±4.44 | 34.73±3.34 * |
| D<sub>15</sub> | 69.28±2.55 | 50.94±3.86 * | 73.62±2.85 | 53.00±2.05 * |
| D<sub>20</sub> | 73.48±2.76 | 57.54±3.34 * | 77.87±2.27 | 56.33±2.19 * |

\*Significant difference ( $p < 0.05$ ) as compared to control

Our study showed that RBCs treated with trypsin and then fixed with glutaraldehyde aggregated in autologous plasma (Fig.3B). At the same time, glutaraldehyde-fixed normal red blood cells (discocytes) do not aggregate in autologous plasma (Fig.3A). Our data are consistent with previous data that aggregation of normal red blood cells fixed with glutaraldehyde is practically absent [2].

**Fig.3.**
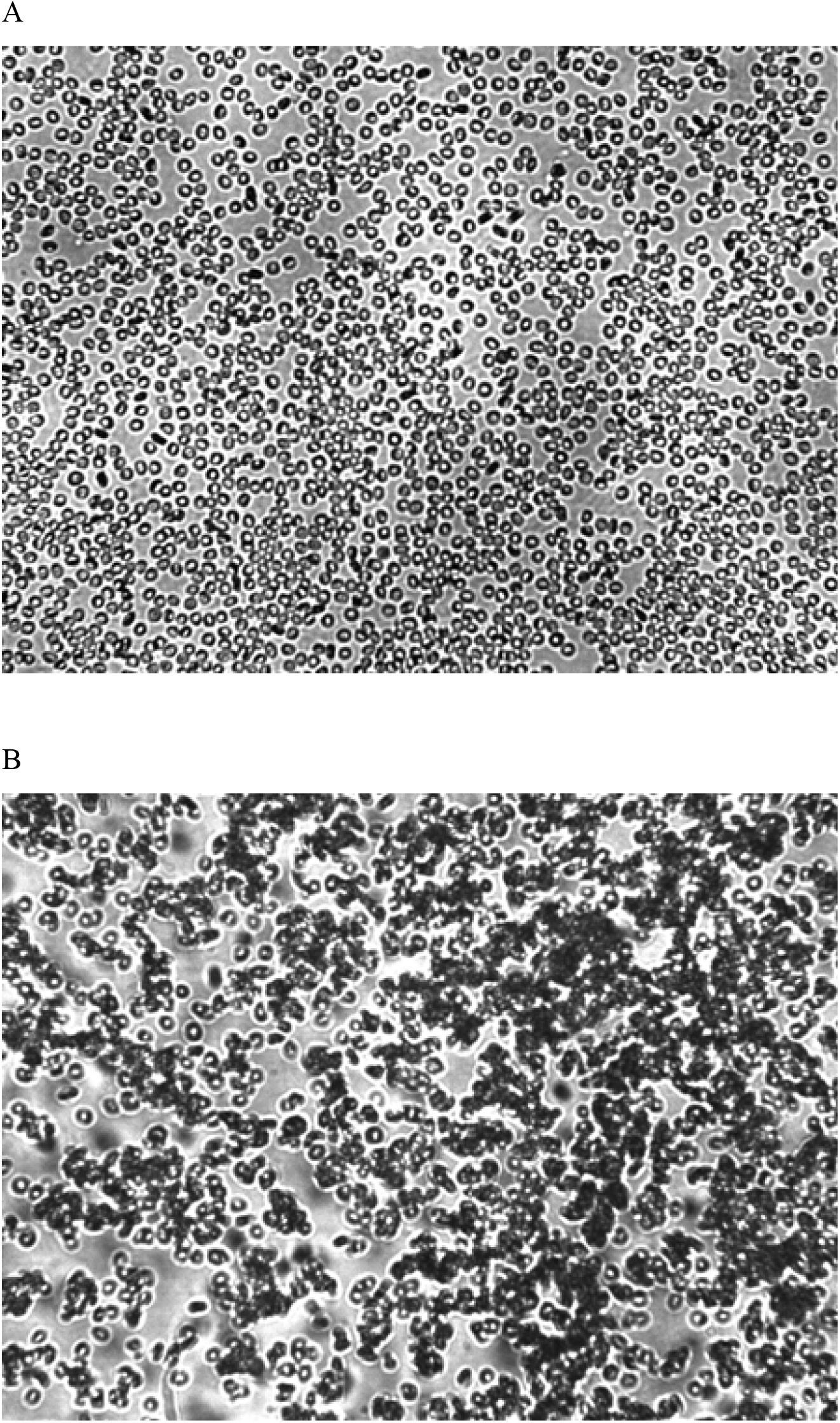
Effects of trypsin - glutaraldehyde treatment on aggregation RBCs: control treated with glutaraldehyde (A) and treated with trypsin and fixed with glutaraldehyde (B) in autologous plasma. Magnification 200x. Treatment with trypsin involved a 30 min incubation of RBCs in the trypsin solution (at hematocrit 10% and at 37^0^C), and then three washes in 10 volumes of buffer. Intact and trypsin treatment RBCs were fixed in a 0.25% solution of glutaraldehyde, and then three washed in 10 volumes of buffer. After that the RBCs were placed in the autologous plasma.

The electrophoretic mobility of normal RBCs in tris buffer averaged − 1.22 ± 0.01μm. s^−1.^V^−1^cm. After trypsin treatment, RBCs showed decrease of electrophoretic mobility to − 0.87±0.01μm. s^−1.^V^−1^cm (ρ <0.001). These results were in agreement with the previous study [3,4,12,14].

## 4. Discussion

The results obtained in this study indicate that trypsin treatment increases RBCs aggregation in autologous plasma and serum. The disaggregation of erythrocytes after trypsin treatment considerably decreased in autologous plasma and serum. The microscopic images of RBCs aggregates indicate the formation of globular (pathologic) structures of aggregates in autologous plasma and serum. Trypsin decrease the surface charge of RBCs. In autologous plasma and serum, the cup shapes of RBCs appear. The control RBCs fixed with glutaraldehyde were not aggregated after their placement in autologous plasma. At the same time, red blood cells pretreated with trypsin and fixed with glutaraldehyde interact with each other in autologous plasma.

In the present research, the aggregation of RBCs was studied in autologous plasma. According to previous research, marked aggregation occurs not only in autologous plasma, but and in autologous serum [23]. Thereupon, we studied RBCs aggregation in not only in autologous plasma, but also in autologous serum. It was found out that trypsin increases RBCs aggregation in autologous serum. Increase in the strength of RBCs aggregates was observed in autologous serum. The morphology of RBC aggregates after trypsin treatment showed the formation of pathologic structures of aggregates in autologous serum.

An analysis of the results obtained in this study showed that the aggregation and aggregate morphology of RBCs after trypsin treatment differ little from the data obtained in patients with diabetic foot disease [20].

Carboxyl groups of sialic acid determine the negative charge of human red blood cells by more than 80% [24]. The integral protein glycophorin A represents 75% of all sialoglycoproteins in the membrane of RBCs [25]. Trypsin has a proteolytic effect only on glycophorin A [26–28]. A decrease in the content of sialic acid was observed in the main membrane of RBCs, glycoprotein glycophorin A, in patients with diabetes [29].

RBCs treated with trypsin showed enhanced susceptibility to cup shape formation by albumin (Mehta 1983). Apparently, the appearance of cup-shaped forms of RBCs in autologous plasma and serum after treatment with trypsin is associated with albumin, autologous plasma and serum. Results indicate an involvement of membrane integral proteins in mediating the shape modulating effects of albumin [30]. Previously shown that the native structure of glycoproteins on the RBCs surface limits normal RBCs aggregation [4]. We have shown that red blood cells fixed with glutaraldehyde after treatment with trypsin aggregate in autologous plasma. This indicates that new sites for binding blood proteins appear on the membranes of RBCs after trypsin treatment.

In conclusion, we can suggest a possible mechanism for increasing the degree of aggregation of red blood cells under the influence of trypsin. Trypsin has a proteolytic effect on glycophorin A. As a result, sialic acid is released, which leads to a decrease in the negative charge. Conformational changes in glycophorin A lead to the appearance of new sites for binding blood proteins. Further formation of protein bridges between neighboring cells leads to aggregation of RBCs.

